# The role of motor and environmental visual rhythms in structuring auditory cortical excitability

**DOI:** 10.1101/2020.04.16.044917

**Authors:** M.N. O’Connell, A. Barczak, T. McGinnis, K. Mackin, T. Mowery, C.E. Schroeder, P. Lakatos

**Author notes:** Corresponding authors: Monica Noelle O’Connell, Ph.D., Translational Neuroscience Division, Nathan S. Kline Institute for Psychiatric Research, Orangeburg, NY 10962, USA, Peter Lakatos, M.D., Ph.D., Translational Neuroscience Division, Nathan S. Kline Institute for Psychiatric Research, Orangeburg, NY 10962, USA.

## Abstract

One of the ways we perceive our external world is through the process of active sensing in which biological sensors (e.g. fingers and eyes) sample the environment utilizing mostly rhythmic motor routines. Previous studies indicate that these motor sampling patterns modulate neuronal excitability in sensory brain regions by entraining brain rhythms, a process termed motor-initiated entrainment. Additionally, rhythms of the external environment, that are independent of internal motor commands, are also capable of entraining rhythmic brain activity. The goal of our study was twofold. First, we aimed to investigate the properties of motor-initiated entrainment in the auditory system using the most prominent motor sampling pattern in primates, eye movements. Second, we wanted to determine whether/how motor-initiated entrainment by eye movements interacts with visual environmental entrainment. By examining laminar profiles of neuronal ensemble activity in the primary auditory cortex of non-human primates, we found that while motor-initiated entrainment has a suppressive, visual environmental entrainment has an enhancive effect. We also found that the two processes are temporally coupled during free viewing, and their temporal relationship ensures that their effect on neuronal ensemble excitability is complementary rather than interfering. Taken together, our results provide strong evidence that motor and sensory systems continuously interact in orchestrating the brain’s rhythmic context for the optimal sampling of our multisensory environment.

## INTRODUCTION

A common tool used by human and non-human animals for the goal dependent exploration of the external world is “active sensing”, a mechanism which employs rhythmic motor initiated sampling patterns in order to create a structured sensory input pattern (Kleinfeld et al., 2006; Schroeder et al., 2010). The likely reason for this is that the brain’s neuronal activity is inherently rhythmic, influencing both the generation of motor activity and the perception of sensory inputs (Buzsáki, 2006), which go hand in hand during active sensing. Previous studies (see (Benedetto et al., 2020) for a review), indicate that rhythmic motor sampling patterns are capable of adjusting neural excitability fluctuations in sensory brain regions to align with the rhythm of the motor production signals (Sperry, 1950). The alignment of rhythmic excitability fluctuations to some other, internal or external rhythmic process is termed neuronal entrainment. Several prior studies have demonstrated that entrainment is a ubiquitous brain mechanism: it is both supramodal and can affect rhythmic brain excitability fluctuations across a range of frequencies (Lakatos et al., 2019). For instance, delta-band oscillatory activity in the whisker barrel cortex of mice has been shown to phase lock to the rate of respiration (1-3Hz) (Ito et al., 2014), and exploratory rhythmic whisking (5-15Hz) results in phase locking of theta/alpha oscillations in both vibrissa primary motor and somatosensory cortices (Ahrens and Kleinfeld, 2004). Rhythmic eye-movements (3-5Hz) have been shown to entrain delta/theta activity resulting in modulation of neuronal excitability in primary visual cortex (Barczak et al., 2019; Ito et al., 2013; Melloni et al., 2009; Rajkai et al., 2008). Lakatos and colleagues (Lakatos et al., 2019) termed these types of entrainment as “entrainment by voluntary self-produced rhythms,” and theorized that a corollary discharge signal from the motor system (Sommer and Wurtz, 2008) is most likely responsible. For clarity, in the current paper we are going to use the term motor-initiated entrainment to describe this type of entrainment, and to distinguish it from “environmental entrainment”.

Environmental entrainment describes a mechanism by which brain rhythms can be entrained by attended or otherwise salient rhythms of the external environment that are independent of internal motor commands and sensory modality (Lakatos et al., 2019; Sameiro-Barbosa and Geiser, 2016). A plethora of studies have shown predictive neuronal excitability modulation through entrainment of low frequency neuronal oscillations by rhythmic auditory and visual stimuli (Barczak et al., 2018; Besle et al., 2011; Cravo et al., 2013; Henry and Obleser, 2012; Lakatos et al., 2008, 2013; Luo and Poeppel, 2007; Mathewson et al., 2012; O’Connell et al., 2014, 2015; Spaak et al., 2014). However, the effect of rhythmic visual stimulation and eye movements on neuronal excitability in A1 is still not known. In general, the effects of motor-initiated vs. environmental entrainment are studied separately, therefore, we decided to investigate their effect on neuronal excitability jointly, as this might reveal important aspects of real-world perceptual processes.

The goals of our current study were to first, confirm that entrainment of neuronal activity occurred in primary auditory cortex (A1) to an environmental rhythmic visual stimulus sequence (i.e. stream of LED flashes). Second, we wanted to investigate the effect of rhythmic motor-initiated visual sampling patterns (i.e. saccades) on neuronal excitability in A1. The reason for investigating these mechanisms in A1 was twofold. First, on a theoretical level, if both types of entrainment co-exist in A1, they would have a profound, so far overlooked effect on audio-visual multisensory interactions. Second, on a practical (signal processing) level, in non-human primates, visual stimulation does not produce large amplitude, evoked type responses, as indexed by transient changes in neuronal firing, at the level of A1 (Kayser et al., 2008; Lakatos et al., 2009). Therefore, we reasoned that A1 might be an ideal location to examine the perhaps conjoint modulatory mechanisms related to eye movements and environmental visual events, as the problem of evoked type, large responses contaminating oscillatory measures is negligible (Obleser et al., 2017).

We analyzed laminar profiles of neuronal ensemble synaptic activity (indexed by current source density (CSD)) and neuronal firing (indexed by multiunit activity (MUA)) recorded via linear array multielectrodes positioned in A1 of awake macaques during two conditions: 1) a rhythmic LED flash stream and 2) in the absence of any stimuli (resting state condition). Eye position was continuously monitored throughout. As expected from earlier work (Lakatos et al., 2009), rhythmic visual stimuli produced entrainment of excitability fluctuations (slow rhythmic fluctuation in phase concentration largely confined to the rate of stimulation, with an accompanying slow fluctuation of neuronal firing), with maximal neuronal firing, signaling high excitability, predictably occurring around LED flash onset. Regarding eye-movements, we found that both in the resting sate and visual stimulation conditions, cortical excitability in A1 was also significantly entrained by saccades. However, as opposed to visual environmental entrainment by rhythmic LED flashes, neuronal ensemble excitability was entrained to its low excitability phase signified by suppressed MUA prior to and around the timing of saccades.

If these two effects, visual entrainment of neuronal ensemble activity to a high excitability state, and entrainment by saccades to a low excitability state, were temporally independent, they would interfere and could cancel each other’s influence. Since the brain tends to strive for efficiency, we examined the temporal relationship of visual stimuli and saccades. We found that saccade probability increased about halfway between two LED flashes, indicating that the brain avoids sensory-motor interference by aligning the timing of motor sampling patterns and sensory inputs in a way that results in complimentary excitability modulation in sensory neuronal ensembles.

## RESULTS

The first goal of our study was to investigate the effect of rhythmic non-auditory environmental inputs (LED flashes) on neuronal ensemble excitability in A1. Second, we wanted to determine if saccading, an internally generated quasi-rhythmic motor sampling pattern, has a similar effect. Additionally, if both modes of entrainment co-occur, how would they interact with one another? The investigation of these “subthreshold”, modulatory mechanisms in A1 is greatly facilitated by the fact that visual stimulation does not result in high amplitude evoked type responses, which could overshadow any subthreshold modulatory influences (Obleser et al., 2017).

### Entrainment by rhythmic environmental stimuli and by saccades in A1

Laminar field potential and concomitant multiunit activity (MUA – indexing spiking activity in local neurons) profiles were obtained with linear array multicontact electrodes (Figure 1A) from 50 A1 sites in 3 awake macaque monkeys under two conditions: 1) during the presentation of a rhythmic (1.8Hz) stream of LED flashes (visual stimulation condition), and 2) in the absence of any stimuli (resting state condition). There was no task required in either condition, and thus monkeys were free to look around anywhere and at any pace. Only one trial block of each condition was recorded in each of the 50 A1 sites. Eye position was continuously monitored using the EyeLink 1000 system (SR Research Ltd). CSD profiles were calculated from field potential profiles and index the location, direction, and density of transmembrane current flow, which is the first-order neuronal response to synaptic input (Nicholson and Freeman, 1975; Schroeder et al., 1998). Figure 1B shows representative laminar CSD and MUA profiles to a stream of LED flashes with an apparent rhythmic, stimulus structure related excitability modulation: across all layers MUA is increasing approaching the onset of the LED flash, while approximately 250ms before and after LED onset – i.e., roughly halfway between LED flashes – there is MUA suppression. This is accompanied by a low amplitude rhythmic CSD fluctuation. In particular, the supragranular layer shows a current source (colored blue, indicating net outward transmembrane current flow) over a current sink (colored red, indicating net inward transmembrane current flow) configuration around LED onset (marked by red arrow). This changes to a sink over source pattern around the time of MUA suppression (marked by black arrows). When compared to auditory responses from the same A1 cortical column site (Fig. 1A), it is apparent that the amplitude of the responses related to the LED stream are much lower (note that the laminar CSD and MUA response profiles are on different scales), and they are more supragranularly weighted. This illustrates that as previously shown for other, non-auditory stimuli, the LED related response in A1 is modulatory (Lakatos et al., 2007, 2009): while there is no abrupt pre-to post-LED flash related change in MUA amplitude (Fig. 2A inset boxplot), and no apparent granular layer CSD activity which would indicate an evoked type response, there is a small rhythmic modulation of local neuronal excitability as indexed by slight CSD and MUA fluctuations, indexing neuronal entrainment (Lakatos et al., 2019).

**Figure 1:**
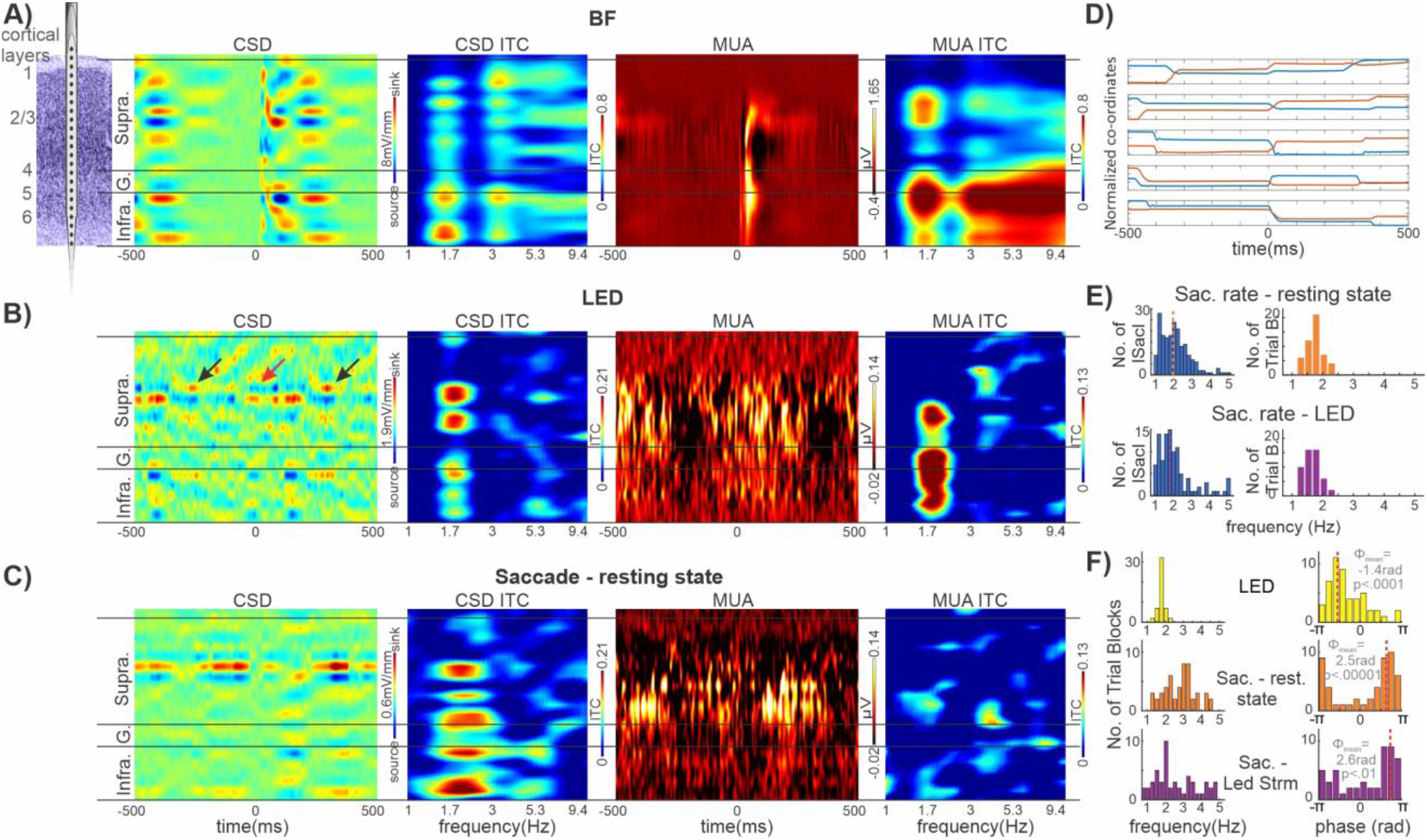
Entrainment by rhythmic environmental stimuli and by saccades in A1. **A)** Schematic of a linear array multielectrode positioned in A1. To the right are representative laminar response profiles of best frequency tone related CSD, CSD inter-trial coherence (ITC), MUA, and MUA ITC. In the ITC plots, frequency is on the x-axis and cortical space is on the y-axis. Higher ITC values mean that the oscillatory phase is more similar across trials while lower values mean that phase values are more random across trials. **B)** Same laminar profiles from the same A1 site as in panel **A** but related to rhythmic LED flashes (1.8Hz repetition rate). **C)** Same laminar profiles and A1 site as A & B but aligned to saccade onset during resting state. **D)** Five examples of horizontal (red) and vertical (blue) eye position traces centered on a detected saccade onset (0ms). **E)** Left bar graphs show representative distributions of intersaccadic intervals (ISacI) rates during a resting state trial block (top) and an LED stream trial block (bottom) from the same experiment. The mean saccadic rate for the two types of trial blocks is represented by vertical dotted lines. Right bar graphs display the mean rate of saccades for all trial blocks during resting state (orange, n=50) and visual stimulation conditions (purple, n=50). **F)** Left column of bar graphs displays the pooled frequencies of the maximum ITC value calculated from the translaminar MUA signal aligned to the timing of LED flashes during LED trial blocks (yellow, N=50), the timing of saccades during resting state blocks (orange, N=50) and the timing of saccades during LED trial blocks (purple, N=50). Right column of histograms show the distribution of the corresponding mean phases. The value of angular mean of the mean phases (marked by red dotted lines), and the rayleigh p-value are inset. Note that LED related ITC and mean phase data (top row) and saccade timing related data (bottom row) are extracted from the same LED trial blocks.

**Figure 2:**
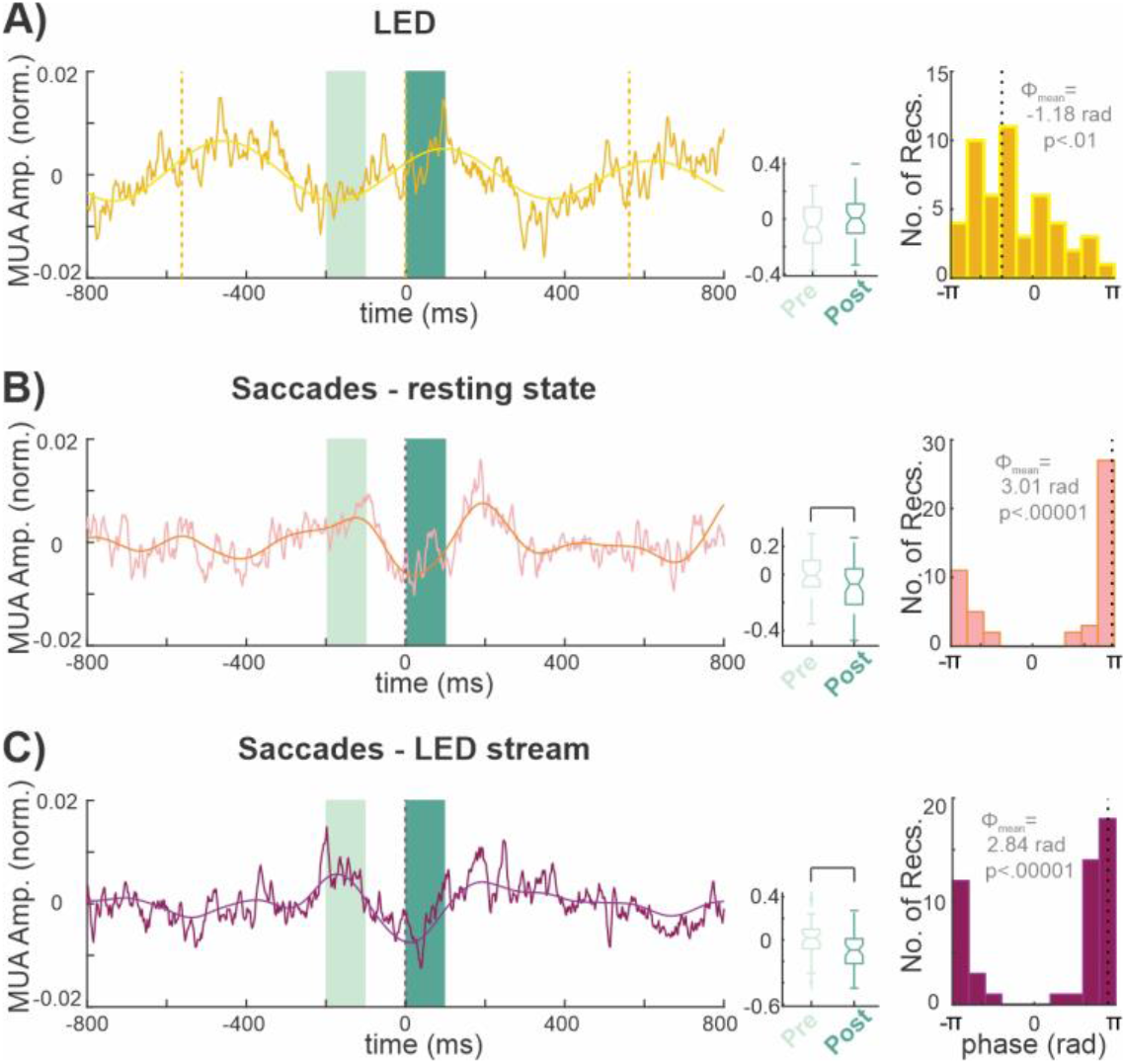
Countersign modulation of excitability by visual environmental stimuli (LED) vs. motor guided sampling of the visual environment (Saccades). **A)** Dark yellow trace shows the rhythmic LED flash related translaminar MUA averaged across all 50 trial blocks. The overlaid bright yellow waveform is the filtered MUA (1.2-2.4Hz). Yellow dotted lines mark LED flash onsets. Shaded light green block marks pre-event (−200 --100ms) while shaded dark green block marks post-event (0 – 100ms) time interval we used to quantify event related MUA modulation. Boxplot to the right shows the pooled normalized MUA amplitudes averaged within the pre-LED flash and post-LED flash onset timeframes. The histogram on the right shows the distribution of mean MUA delta phases at LED flash onset calculated from the bandpass filtered MUA data. The value of angular mean of the mean phases (marked by dotted lines), and the rayleigh p-value are inset. **B)** Pink trace shows translaminar MUA aligned to saccade onset and averaged across all 50 trial blocks during the resting state condition. The overlaid orange waveform is the filtered MUA (1-5Hz). Boxplot to the right shows the pooled MUA amplitudes averaged within the pre-saccade and post-saccade onset timeframes. Bracket marks a significant difference (Wilcoxon signed rank, p < 0.01). The histogram to the right shows the distribution of mean MUA delta/theta phases at saccade onset from the bandpass filtered MUA data. Dotted line marks angular mean of the mean phases. **C)** Same saccade timing related data as in B, but for trial blocks during which the rhythmic LED stream was presented (same trial blocks as panel A).

To provide further evidence for this notion, we investigated the phase coherence of oscillatory activity across trials (indexed by intertrial coherence – ITC) (Lakatos et al., 2005a, 2007). ITC indexes the frequency specific phase consistency of neuronal activity across trials, which, in the case of entrainment should be non-random, i.e. pooled around a specific phase. ITC values vary between 0 and 1, and higher values indicate stronger pooling of phases. As the laminar ITC profiles to the right of the CSD and MUA profiles show, auditory responses (Fig. 1A) are characterized by “broadband” high ITC values, typical of an evoked response (Lakatos et al., 2007, 2009). As opposed to this, LED stream related neuronal activity has only one prominent peak around 1.8Hz, which corresponds to the presentation rate of LED flashes (Fig. 1B). This was the case for all 50 LED stream trial blocks. To summarize, the modulatory type responses to LED flashes coupled with significant phase similarity (pooling) across trials at the frequency of stimulus presentation indicate oscillatory entrainment (Lakatos et al., 2019; Obleser and Kayser, 2019).

Next we examined the effect of saccades on auditory cortical activity, reasoning that since our results show that visual inputs modulate A1 excitability, saccade related inputs, which has been shown to modulate excitability in primary visual cortex (Barczak et al., 2019; Rajkai et al., 2008), should as well. If entrainment could be brought about by the quasi-rhythmic sequence of saccades (Amit et al., 2017; Henderson, 2003), we expected that the neuronal activity would be entrained to the same excitability phase as in the case of the LED stream, as both saccades and LED flashes are associated with a post-event volley of visual input. We tracked eye movements at 500Hz resolution using an infrared system (see Methods). Figure 1D shows representative saccade triggered (positioned at 0ms) horizontal and vertical eye-position traces. Note that in all 5 examples, there are “flanking saccades” occurring approximately 400ms prior to and following the saccade the traces are aligned to. This is due to the quasi-rhythmic nature of saccades, which was demonstrated by numerous human and non-human primate studies (Barczak et al., 2019; Hoffman et al., 2013; Maldonado et al., 2008; Rayner, 1998) and is illustrated by Figure 1E. The pooled mean rate of saccades we detected (Fig. 1E right) is similar for both resting state and LED trial blocks (mean saccade rate is 1.76Hz (standard deviation (SD) = 0.25Hz) and 1.69Hz (SD = 0.24Hz) respectively), which is slightly lower than in previous non-human primate studies (Barczak et al., 2019; Berg et al., 2009; Ito et al., 2011), likely due to the lack of a visual task and the stricter criteria used for saccade detection (see Methods). Figure 1C shows representative laminar CSD and MUA profiles aligned to saccade onsets (n=325 saccades) recorded during the resting state condition from the same A1 site as in A and B. Similar to the profiles related to the LED stream, there is a near baseline modulation of the MUA and a relatively low amplitude CSD fluctuation again focused in the supragranular layers. This is accompanied by a prominent ITC peak around 2Hz (mean rate of saccades in this particular trial block) in the CSD ITC plot and to a lesser extent in the MUA ITC plot, which indicates that – in lieu of evoked type activity – neuronal oscillations did become aligned to the timing of saccades. In other words, similar to a rhythmic stream of LED flashes, a quasi-rhythmic series of saccades can also entrain neuronal activity in A1.

### Countersign modulation of A1 excitability by rhythmic LED flashes vs. Saccades

Since neuronal oscillations reflect rhythmic fluctuations of excitability in neuronal ensembles (Buzsáki, 2006), next we wanted to see whether oscillatory activity was being entrained to the high or low excitability phase in each of these cases. Since the MUA signal has a straightforward relationship to excitability, and the excitability of all cortical layers tends to oscillate synchronously within a given A1 column (Lakatos et al., 2013; O’Connell et al., 2011), we used translaminar MUA (i.e., averaged across all cortical layers) to estimate “excitability phases”: high amplitude MUA corresponds to high, while low amplitude MUA corresponds to low excitability of neuronal ensembles in A1 (Lakatos et al., 2005a). Given that the frequency of saccades varied across trial blocks (Fig. 1E, right columns), we initially checked the degree of oscillatory phase-similarity (i.e. ITC), and mean phases in the 1-5 Hz frequency range. Left columns in Figure 1F show the pooled frequencies of maximum ITC values determined at onset of LED flashes and at saccade onset during the two conditions. As expected, LED stimulus stream related ITC values are tightly grouped around 1.8 Hz (which is the stimulus presentation rate), while saccade related ITC values are more widely distributed, both during resting state and LED stream conditions. When we looked at the matching mean phases (at the peak of ITC), we found a significant bias for each of the three groups (Rayleigh’s uniformity tests p < 0.01, Fig. 1F, right column). We also found that the mean phases related to LED flashes and those related to saccade onsets are significantly different. While pooled delta phases calculated at LED flash onset are significantly grouped (Rayleigh’s uniformity tests, p = 0.3 x 10^-4^) before the positive peak (0 rad) of the MUA oscillation which corresponds to the high-excitability phase of the MUA oscillation, pooled delta/theta phases calculated at saccade onsets are also significantly grouped (Rayleigh’s uniformity tests, during resting state p = 0.13 x 10^-5^, and during visual stream p =0.0012) but around the opposite phase (±pi rad), which corresponds to the low-excitability phase of the MUA oscillation.

To verify this unexpected, counterphase entrainment by LED flashes and saccades, we band-pass filtered the translaminar MUA in the 1.2-2.4Hz and 1-5Hz frequency bands respectively, and then calculated the phase of the resulting signal at flash or saccade onset using the Hilbert transform. We used these particular frequency bands as they spanned the frequency range of maximum ITC values in Figure 1F (left column). The yellow histogram in Figure 2A displays the pooled mean delta phase distribution of all trial blocks associated with the LED flashes, which shows a very similar pattern as the wavelet results (Fig. 1F). There is a significant non-random distribution of mean phases (Rayleigh’s uniformity tests, p =0.004) around the high excitability phase, as the grand average translaminar MUA is approaching its positive peak (indexing a high incidental neuronal firing rate) at the time of the LED flashes (Fig. 2A, yellow waveform). For saccade-related activity during resting state, pooled mean delta/theta MUA phases (Fig. 2B) are also significantly grouped (Rayleigh’s uniformity tests, p = 0.9 x10^-14^), but around the opposite, low cortical excitability (indexing a low incidental neuronal firing rate). In fact, there appears to be a clear MUA suppression around saccade onset (0ms) in the corresponding grand average translaminar MUA (Fig. 2B, pink waveform). Saccade related excitability phase results during LED flashes (Rayleigh’s uniformity tests, p = 0.1 x 10^-13^) were almost identical to those during resting state condition (Fig. 2C). The pooled LED related mean phase distributions were significantly different from saccade related phase distributions during both resting state (Fisher’s test for the equality of circular means, p =0.3 x 10^-7^), and during visual stimulation (Fisher’s test for the equality of circular means, p = 0. 3 x 10^-8^). Saccade related mean phase distributions during resting state and visual stimulation conditions were not significantly different (Fisher’s test for the equality of circular means, p = 0.06).

To summarize our results thus far, while rhythmic environmental visual stimuli and quasi-rhythmic motor guided visual sampling both entrain A1 neuronal activity, they do so to opposite neuronal ensemble excitability phases. To quantify this in yet another way, we compared pre- (–200 to −100ms) and post-event (0-100ms) MUA amplitudes. We found that there was only a significant difference between pre- and post-saccade MUA amplitudes (Wilcoxon signed rank, p = 0.0016 during resting state condition, and Wilcoxon signed rank, p = 0.00004 during visual stimulation condition) (boxplots in Fig. 2B & C). While there was a trend towards larger post-LED flash translaminar MUA amplitude, it was not significant (Wilcoxon signed rank, p = 0.10).

### Long- and short-timescale interaction of saccades and environmental visual stimuli

So far, we have demonstrated that environmental visual stimuli and internally generated motor actions controlling eye movements aimed at sampling the visual environment are both capable of entraining neuronal activity in A1. If these influences would operate independently in time their effects would vary randomly between synergy and interference. We thus examined how the two types of entrainment described above (environmental vs. motor-initiated entrainment) interact on long- (second) and shorter (sub-second) time scales. For this analysis, we focused on trial blocks during which LED flashes were presented.

Our first question was: do delta/theta oscillations entrain to LED flashes and saccades simultaneously, or does this vary by trial block, possibly indicating different brain states and/or information sampling strategies. To answer this, we determined the percentage of trial blocks with significant entrainment (Rayleigh’s uniformity tests, p < 0.01) to either the LED flashes only, saccades only, or to both events (comprising three “entrainment mode” groups), in at least one laminar recording location (i.e. a linear array electrode contact) (Fig. 3A-B).

**Figure 3:**
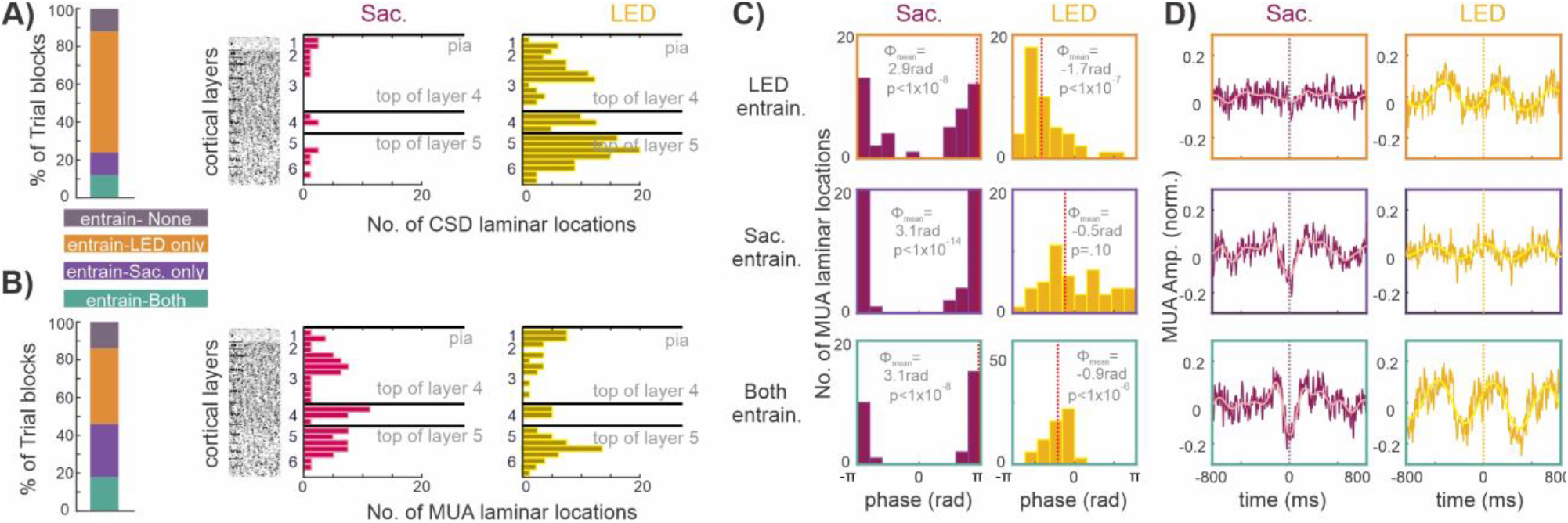
Spatiotemporal properties of A1 modulation elicited by LED flashes and Saccades. **A)** Bar shows *%* of LED trial blocks where at least 1 laminar recording location in which CSD measures were derived, significantly entrained to either the LED flashes (orange), saccades (purple), both (green) or none (grey) (Rayleigh p < 0.01), constituting 3 “entrainment mode” groups. Bar graphs to the right display the number of CSD recording locations showing significant entrainment (saccade related N= 17, LED related N= 144) relative to top of cortex (pia), top of the granular (layer 4) and top of the infragranular (layer 5) layers, all marked by black horizontal lines. For reference, we show an A1 slice stained for Nissl substance (reproduced from Fig. 4, (Hackett et al., 2001)). **B)** Same as in A but for MUA laminar recording locations showing significant entrainment: saccade related N=76, LED related N=68. **C)** Histograms show the distribution of mean MUA phases related to saccades (maroon) and LED flashes (yellow) calculated at MUA laminar recording locations which significantly entrained to LED (orange outline), saccades (purple outline) or both (green outline). The value of angular mean of the mean phases (marked by red dotted lines), and the rayleigh p-value are inset. **D)** MUA recording locations aligned to saccades (maroon) and LED flashes (yellow) pooled by “entrainment modes”. The overlaid waveforms display the average filtered MUA (pink: 1-5Hz, bright yellow: 1.2-2.4Hz).

This analysis also gave us the opportunity to examine the laminar distributions of all electrode contact locations showing significant entrainment to saccades vs. LED flashes, which is described in the next three paragraphs. To enhance this aspect of the analysis, besides laminar MUA, we also included laminar CSD data. By laminar CSD recording locations we mean the electrode contact recording site at which CSD was calculated (Freeman and Nicholson, 1975; Mitzdorf, 1985). The left bar graph in Figure 3A shows CSD entrainment occurred in most trial blocks (64%, N=32) to the LED stream only (environmental entrainment), while 12% (N=6) of trial blocks showed CSD entrainment to saccades only. Additionally, 12% (N=6) showed CSD entrainment to both LED flashes and saccades, and 12% (N=6) failed to show CSD entrainment to either event type. Right bar plots in Figure 3A display the distance of the CSD recording locations which entrained to saccades and LED stream, from the pial surface of A1 (top black horizontal line), relative to the top of the granular (4, middle black line) and top of the infragranular (layer 5, bottom black line) layers. We found that more CSD laminar locations showed entrainment to the LED stream (23%, N=144/640 laminar locations across the 38 trial blocks) than to saccades (8%, N=17/221 laminar locations across the 12 trial blocks). While all cortical layers contain some CSD laminar locations entrained by the LED stream, the distribution appears to be slightly more biased to upper layer 3 and layer 5 (Fig. 3A, yellow bar plot). Due to the low number of laminar CSD locations entraining to saccades, it is difficult to see a laminar pattern but there is a marginal bias to layer 1 (Fig. 3A, pink bar plot).

While admittedly MUA is somewhat volume conducted (Kajikawa and Schroeder, 2015), and stimulus related/modulatory MUA effects also depend on the amount of baseline firing rates in different layers (Petersen and Crochet, 2013), we conducted the same analysis as for the CSD data (Fig. 3B). Similarly, more trial blocks (40%, N=20) showed MUA entrainment in at least one laminar location, to the LED stream only (environmental entrainment). 28% (N=14) of trial blocks showed MUA entrainment to saccades only, whereas 18% (N=9) showed MUA entrainment to both LED flashes and saccades, and 14% (N=7) failed to show MUA entrainment to either event (Fig. 3B, left bar graph). We found that a few more MUA laminar locations entrained to saccades (19%, N=76/403 laminar locations across the 23 trial blocks) than to the LED stream (14%, N=68/485 laminar locations across the 29 trial blocks). Qualitatively, we found that the distributions of laminar locations where MUA entrained to saccades vs. LED flashes were similar in that every layer showed entrainment, however locations showing MUA entrainment to LED flashes appear to be slightly more biased to layers 1 and 5 (Fig. 3B, right bar plots).

The above results (Fig. 3A & B, right bar plots) indicate that there is an apparent difference in the capability of saccades vs. LED stream to align CSD vs. MUA. Therefore, we tested the strength of alignment of CSD and MUA to saccades vs. LED stream, which revealed a strikingly opposite effect. We found in the case of saccade related alignment, MUA ITC values calculated at all individual laminar recording locations were significantly larger than CSD ITC values calculated at the same locations (Wilcoxon signed rank, p = 0.7 x 10^-19^). However, in the case of LED stream related alignment, CSD ITC values were significantly greater than MUA ITC values (Wilcoxon signed rank, p = 0.5 x 10^-4^). The differential influence of saccades vs. LED on neuronal firing (MUA) and transmembrane currents (CSD) might indicate distinct circuit mechanisms.

Next, within each of the three “entrainment mode” groups (LED, saccade, and both) we compared the saccade and LED flash related MUA mean entrainment phases of individual laminar recording locations (Fig. 3C). We found that in general, variation in mean delta/theta phases did not appear related to “entrainment mode”. The only exception is that mean LED related phases were not pooled significantly in the “entrained to saccades” group. Figure 3D displays the entrainment mode specific grand average of the corresponding MUA traces from which the phases were calculated.

To summarize this section to date, we found that in some trial blocks, both saccades and environmental visual stimuli entrain auditory cortical oscillations (e.g. Fig. 3A & B). However, these effects occur when evaluated for all stimuli/saccades occurring across an entire trial block. Thus, we wondered if within a trial block whether periods of entrainment by LED stimuli vs. saccades “co-exist” or rather fluctuate (long-timescale interaction), similar to what has been demonstrated in A1 across different stimulus modalities (Lakatos et al., 2016). To test this, we calculated the phases of the bandpass filtered translaminar MUA at saccade and LED flash onsets in 10 second windows and calculated ITC (moving ITC). The step size of the moving ITC window was 1 second (resulting in a 1 Hz “sampling rate”). As the representative examples in Figure 4A show, the moving ITC measure related to LED flashes (yellow traces) and saccades (maroon traces) varies robustly across a trial block, indicating at times stronger environmental vs. motor-initiated entrainment and vice versa. To quantify this, we next computed the correlation between saccade related and LED related moving ITC. We found a significant negative correlation in 10 of the 50 (20%) LED stream trial blocks (Spearman’s correlation, p < .05), signaling that when moving ITC related to one event is high it is low for the other event, and vice versa. Only 3 of the LED stream trial blocks (6%) showed a significant positive correlation (Spearman’s correlation, p < .05), indicating that moving ITC related to both saccade and LED co-varies together across the entire trial block, while the remaining 37 LED trial blocks (74%) failed to show a significant correlation.

**Figure 4:**
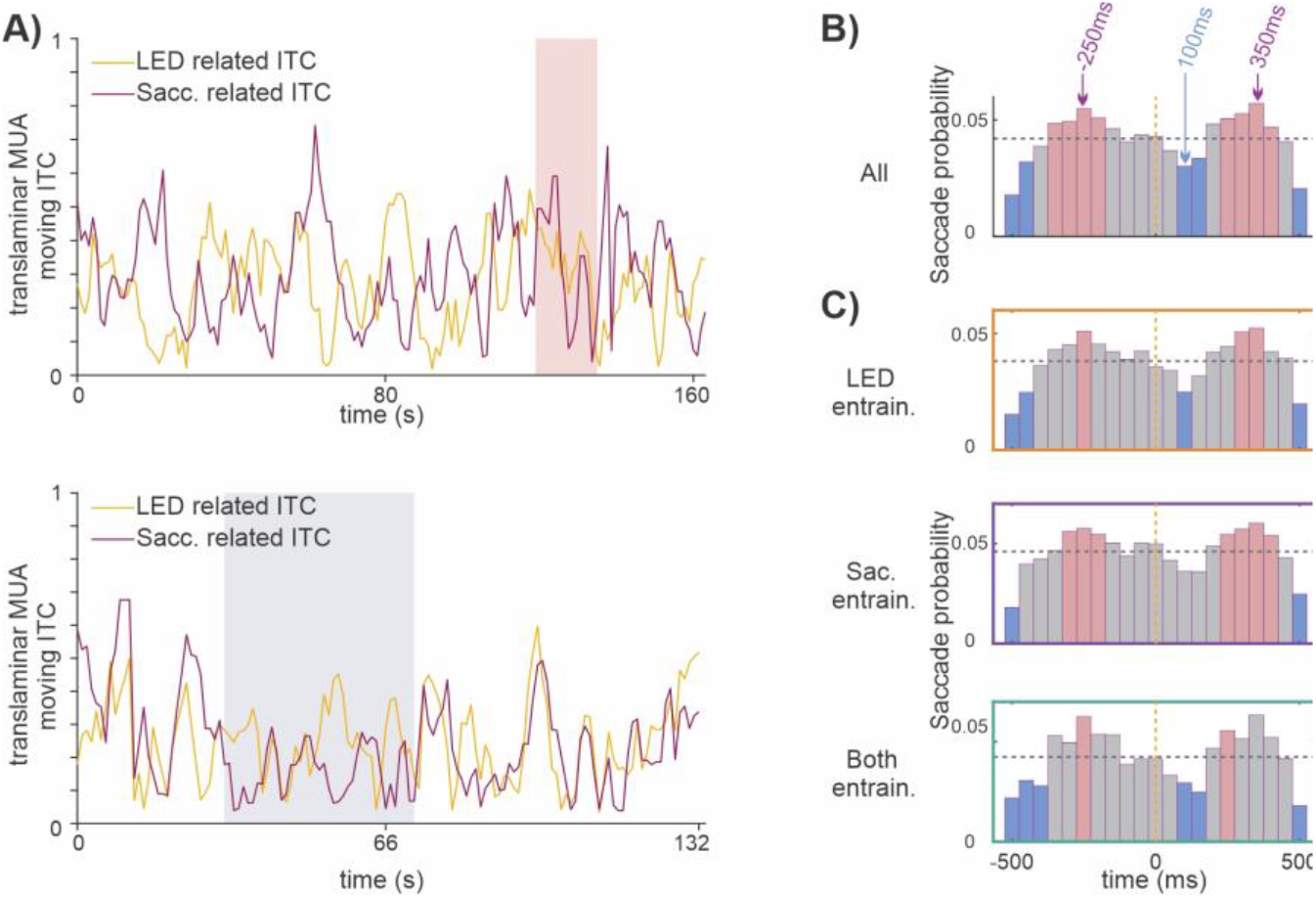
Countersign fluctuation of environmental vs. motor-initiated entrainment, and temporal relationship of saccades and rhythmic LED flashes. **A)** Top: “Moving average” cortical MUA ITC values from a representative trial block showing an overall significant negative correlation (Spearman’s correlation, p < .05) between saccade (maroon) and LED related entrainment (yellow). Phase at the time of LED flashes/saccades was extracted from data bandpass filtered between 1.2-2.4Hz and 1-5Hz respectively. Shaded pink block marks a time period where a significant positive correlation occurred (Spearman’s correlation, p < .05). Bottom: Representative traces from a different trial block showing an overall significant positive correlation (Spearman’s correlation, p < .0001) between saccade and LED related entrainment. Shaded blue block marks a time period where there was significant negative correlation (Spearman’s correlation, p < .05). **B)** Bar graphs show the distribution of saccade probability relative to LED onset (0ms) pooled across all 50 LED trial blocks. Pink bars denote time intervals in which saccades occurred significantly more often (Wilcoxon signed rank, p < 0.05), while blue bars denote when saccades occurred significantly less often (Wilcoxon signed rank, p < 0.05) than the average (marked by horizontal dotted grey line). **C)** Same as in B but for trial blocks pooled by MUA “entrainment modes” (see Fig. 3B).

Somewhat puzzled by this finding, we examined moving ITC correlations in all LED stream trial blocks on a temporally more fine-grained scale. Examining again the representative examples in Figure 4A might possibly explain this finding: while the traces in the top plot show an overall significant negative correlation (r = −0.18, Spearman’s correlation, p =.02), the pink block marks a time period where the moving ITC measures related to saccade and LED show a significant positive correlation (r = 0.76, Spearman’s correlation, p =.0005). Likewise, whereas the bottom moving ITC traces show an overall significant positive correlation (r = 0.37, Spearman’s correlation, p = .00001), the blue block marks a time period where the moving ITC measures related to saccade and LED show a significant negative correlation (r = −0.34, Spearman’s correlation, p =.02). Many of the trial blocks showed a similar pattern. This implies that during an entire trial block there are times where entrainment is exclusive (i.e. A1 is entrained by saccades or LED flashes independently), and other periods where both events seemingly simultaneously entrain neuronal oscillations.

This was a somewhat unexpected finding, as previous studies seem to indicate that entrainment is exclusive to a given modality/feature in A1 if entraining events are temporally independent (Lakatos et al., 2009, 2016). Therefore, we wanted to examine whether the temporal relationship of saccades and rhythmic LED flashes is indeed independent on a short-timescale. As the pooled distribution of saccade probability for all trial blocks show (Fig. 4B) there is an increase in saccade probability about 250ms before and after an LED flash; approximately halfway between two LED flashes. This is accompanied by significantly decreased saccade probability following flashes. When the LED trial blocks are grouped within the “entrainment mode” groups as determined by MUA (see Fig. 3B), the same pattern is evident to a varying degree (Fig. 4C). This indicates that there is a strong short-timescale temporal relationship between the sampling of the visual environment and environmental visual stimuli.

## DISCUSSION

The present study found that both rhythmic environmental stimuli, such as LED flashes, and quasi-rhythmic motor sampling patterns, such as saccades, are capable of entraining rhythmic neuronal excitability fluctuations in A1. Surprisingly, while the LED stream entrained neuronal oscillations to their high excitability phases such that MUA was at its peak around the time when LED flashes occurred (via environmental entrainment), saccades entrained neuronal oscillations to their low excitability phases (via motor-initiated entrainment), which resulted in MUA suppression around saccade onset (Fig. 2). We also found that the two types of entrainment could co-occur in some trial blocks (Fig. 3). However, when we examined this on a longer timescale, we found that within trial blocks there were periods where entrainment was exclusive (i.e. A1 activity is either entrained by saccades *or* LED flashes, but not by both), and other periods during which neuronal activity was aligned to both internal and external events. This latter finding can only be explained if the timing of visual events and saccades is non-independent, i.e. visual inputs entrain saccade generating mechanisms. We indeed found that there was a robust short-timescale temporal relationship between environmental visual stimuli and the sampling of the visual environment, in that saccade probability significantly increased about halfway between two LED flashes and significantly decreased following flashes (Fig. 4B). Since we did not require the animals to perform a task, there is a possibility that based on task demands saccades and environmental rhythms can be decoupled (e.g. during visual search vs. deviant stimulus detection). However, as our results indicate, one of the brains major strategies is to avoid interference by coupling sensory and motor systems most of the time.

### Relevance of temporal coupling of eye movements and environmental visual stimuli

Our findings indicate that saccades are not coincident with LED flashes but do however synchronize their occurrence to these so that they fall reliably about halfway between two flashes in the rhythmic “LED cycle”. This is consistent with results of previous studies that showed saccadic synchronization to rhythmic external stimuli (Batten and Smith, 2018; Joiner et al., 2007; Takeya et al., 2017). While it has been known for a long time that relevant or high intensity sensory stimuli result in reflexive, or reactive saccades (Joiner et al., 2007; Shelhamer and Joiner, 2003), more recent studies (Hayhoe et al., 2012; Henderson, 2017), demonstrated that saccades can also be predictive, i.e. occur in anticipation of stimuli.

With any rhythmic stimulation, like the one we used, the question of whether saccadic timing is reactive or predictive is a valid one, as both post- and pre-stimulus timing is “fixed”. While the median saccadic reaction time to an LED stimulus for regular saccades is 130-180ms in non-human primates (Dorris and Munoz, 1998; Fischer, 1986; Paré and Munoz, 1996), we actually found a decrease in saccade probability in this post-stimulus timeframe (Fig. 4B). This might indicate that the saccade timing observed by us, approximately 250ms before LED flashes is predictive, i.e. saccades occur in anticipation of a temporally predictable environmental event. In addition, a series of studies by Joiner and colleagues have shown that when rhythmic stimuli are presented around 2Hz (similar to the current study) saccades are predictive, compared to stimuli presented at lower rates which result in reactive saccades (Joiner et al., 2007; Shelhamer and Joiner, 2003; Yuval-Greenberg and Deouell, 2011). Future studies using quasi-rhythmic visual stimulus streams, that have some jitter in time between single stimuli instead of strictly isochronous sequences could verify this notion.

Whatever the case (predictive or reactive saccade timing), our results indicate that the brain “binds” environmental and motor events so that their countersign effect on the excitability of neuronal ensembles does not interfere with each other. Even though our motor actions are governed by behavioral goals, evidence suggests that the brain governs our actions so that they (e.g. rhythmic sampling patterns) are guided by the timing that inherently exists in the internal and external environment (Lakatos et al., 2019).

### The circuitry of saccade related entrainment in A1

Strikingly, while delta phase alignment related to the LED stream was prevalent in the CSD, in relation to saccade timing we found the reverse: more locations across layers showed MUA rather than CSD entrainment (Fig. 3A-B). This indicates that while as previously demonstrated, multisensory (i.e. LED related) alignment of neuronal activity affects CSD more, saccade related effects influence neuronal firing (MUA) more than transmembrane current flow (CSD). Since the CSD signal is dependent on cell geometry more than MUA, it is likely that motor related inputs affect different cortical neuronal networks than LED flash related inputs, which in turn might indicate a different function for motor vs. environmental modulation of neuronal excitability. We speculate that there are four hypothetical circuits underlying saccade related entrainment in A1, that could be verified by future anatomical and neuromodulation studies.

The *first* and most obvious would be direct projections from motor cortex transmitting a corollary discharge signal related to the production of eye movements. Anatomical studies in rodents have identified direct projections from both primary and secondary motor areas to A1 (da Costa et al., 2017; Nelson et al., 2013), whose activation suppresses neuronal activity in auditory cortex (Nelson et al., 2013; Schneider et al., 2014, 2018). These motor areas most densely innervate the supragranular and infragranular layers of auditory cortex (Nelson et al., 2013). However, these connections have not been observed in primates. As seen in the current and many previous studies, phase reset and entertainment of ongoing oscillations is most prevalent in extragranular layers (Lakatos et al., 2005a, 2007, 2009; O’Connell et al., 2011), and modulatory inputs targeting these layers have long thought to cause phase reset, and if stimuli are rhythmic, entrainment (Barczak et al., 2019; Lakatos et al., 2009; O’Connell et al., 2011, 2014). Besides anatomical studies, recent magnetoencephalography studies investigating predictions in auditory scenes in humans found significant effective connectivity from motor cortex toward auditory areas (Abbasi and Gross, 2020; Morillon and Baillet, 2017). Additionally, besides direct projections, an indirect source of motor influences on the auditory system could stem from the cochlea, as a recent study reported oscillations of the eardrum coinciding with saccade onset in the absence of sound (Gruters et al., 2018). The authors hypothesized that the eardrum movement is due to a copy of the motor command that generates saccades. It is possible that corollary discharge signals originating from motor cortex affect the activity of the middle ear muscles and neuronal activity of A1 in parallel. It is also conceivable that the movement of the eardrums alone modulates oscillatory activity in A1 through the non-lemniscal auditory matrix pathway, which also targets the superficial layers (Jones, 1998; Molinari et al., 1995).

The *second* possible circuit, not involving the motor cortex, could be based on a superior colliculus -> pulvinar -> A1 routing of saccade related corollary discharge. It is well established that the superior colliculus is involved in the generation of saccades (Krauzlis et al., 2013; Moschovakis et al., 1996; Olivier et al., 1993; Robinson and Wurtz, 1976). Particular neurons in the deeper, or “motor” layers generate bursts of action potentials that command saccades (Wurtz and Goldberg, 1972), and these deep, non-retinorecipent layers also send projections to the pulvinar (Baldwin et al., 2011). The pulvinar has been shown to be active during saccades but not prior to (Robinson et al., 1986), indicating its engagement by the saccadic network independent of visual inputs. The pulvinar is the largest multimodal nucleus of the thalamus and possesses extensive reciprocal cortical connections with the upper layers of sensory cortices including A1 (Homman-Ludiye and Bourne, 2019; Homman-Ludiye et al., 2019). In support of this possible circuit for saccade related MUA suppression, a recent study has shown that activation of pulvinar projections targeting neurons in the superficial layers of A1, especially layer 1, leads to a suppression of A1 neuronal activity (Chou et al., 2020).

A *third* feasible source for saccade related modulation of the auditory system and the temporal alignment of predictable environmental stimuli and rhythmic motor sampling are the basal ganglia. Studies have shown that previous experience creates stimulus response associations through the basal ganglia circuit that promote the synchronization of motor eye fields (saccades), auditory, and visual sensory systems through orientation responses (Boussaoud and Joseph, 1985; Hikosaka et al., 1989, 1993; Joseph and Boussaoud, 1985). The cerebral cortex projects to the dorsal and ventral striatum which send projections to the output nuclei of the basal ganglia which ultimately project back to the sensory cortex via the thalamus, forming a closed loop (Alexander et al., 1986; Lim et al., 2014). In this way, internally generated sensorimotor responses to previous experience (like our rhythmic light stimulus sequence) and externally generated sensorimotor responses to ongoing stimuli are modulated by feedforward (motor internal) and feedback (sensory external) input to basal ganglia circuits producing anticipatory effects on neural activity.

Finally, saccade related entrainment in A1 could be orchestrated by a *fourth* possible source, in this case sensory as opposed to motor: the retinal input volley related to each fixation following most saccades. It has previously been shown that attention to visual stimuli can cause phase reset of ongoing oscillations in A1 (Lakatos et al., 2009). In theory the same mechanism could be responsible for the saccade related entrainment seen in the current study. A possible pathway for the inputs responsible for this type of entrainment could be via the multisensory part of the medial geniculate nucleus, the medial or magnocellular division (MGNm), as it sends thalamocortical projections to layers 1 and 3 of A1 (Bartlett, 2013; Huang and Winer, 2000; Mitani et al., 1987). MGNm receives inputs from all layers of the superior colliculus including the lower layers which are multimodal (Benevento and Fallon, 1975; Linke, 1999). However, our results indicate that this circuit mechanism is the least likely one. The reason for this is that if post-saccadic visual inputs (i.e. visual input volley) would be responsible for saccade related modulation of A1, they should entrain A1 neuronal ensembles to their high excitability phases like environmental stimulus (i.e. LED) related visual inputs. In contrary, we found that saccades and environmental visual inputs entrain neuronal excitability to opposing phases in A1.

### Motor related suppression of Sensory Cortices

Apart from the well documented saccadic related suppression in visual cortex (for a review see (Gegenfurtner, 2016)), a plethora of studies have shown in both humans and animals that self-produced motor activity (e.g. button-pressing, vocalizing, walking, etc.) suppress neural activity in auditory cortex (Aliu et al., 2009; Eliades and Wang, 2003; Flinker et al., 2010; Houde et al., 2002; Schneider et al., 2014). In fact Eliades and Wang showed predictive suppression of neuronal activity in the auditory cortex of marmosets prior to and during self-initiated vocalizations (Eliades and Wang, 2003). This phenomenon is not restricted to the auditory cortex, as responses in somatosensory cortex to self-produced tactile stimuli are attenuated compared to the same stimuli but when externally generated (Blakemore et al., 1998). The results of the current study are in line with and extend these findings, since we show that self-generated quasi-rhythmic motor acts result in a rhythmic suppression of auditory cortical activity. While the exact relevance of this mechanism in the case of eye-movements is unknown, we speculate that during saccading, there should be no associated auditory stimulus, as saccades are “silent”. Nonetheless, if random auditory stimuli occur concurrent with saccades, these would be distracting to the brain’s goal, which is to resolve the visual environment. Therefore, it is beneficial to suppress them around the timing of saccades.

While there is evidence that rhythmic motor activity enhances sensory perception (Morillon et al., 2014), to our knowledge, there is no direct evidence for motor related enhancement of excitability in sensory cortical regions. Our results lead us to think that by aligning motor activity to sensory inputs, the brain reduces interference from motor related suppression, which, together with the rebound in excitability following motor inputs indirectly enhances sensory processing. In fact there already appears to be some evidence for this in the visual system (Barczak et al., 2019).

## Conclusions

Our study provides evidence that both visual environmental rhythmic stimuli and quasi-rhythmic eye movements are capable of entraining neuronal activity in A1. While environmental visual stimuli entrain oscillations to their high excitability phases, saccades entrain them to their low excitability phases. We propose that this is due to the brain’s opposing intentions in the two cases: in the first, environmental case, an enhancement of co-occurring auditory inputs is beneficial in order to facilitate multisensory integration, while in the second, motor controlled sensory sampling circumstance, it is advantageous to avoid interference of inputs related to random environmental stimuli.

## MATERIALS AND METHODS

### Subjects

In the present study, we analysed electrophysiological data recorded during 50 penetrations of area A1 of the auditory cortex of 1 male and 2 female rhesus macaques (17, 31 and 2 penetrations respectively) weighing 5-8 kg, who had been prepared surgically for chronic awake electrophysiological recordings. Prior to surgery, each animal was adapted to a custom fitted primate chair and to the recording chamber. All procedures were approved in advance by the Animal Care and Use Committee of the Nathan Kline Institute.

### Surgery

Preparation of subjects for chronic awake intracortical recording was performed using aseptic techniques, under general anesthesia, as described previously (Schroeder et al., 1998). The tissue overlying the calvarium was resected and appropriate portions of the cranium were removed. The neocortex and overlying dura were left intact. To provide access to the brain and to promote an orderly pattern of sampling across the surface of the auditory areas, Polyetheretherketone (PEEK) recording chambers (Rogue Research Inc.) were positioned normal to the cortical surface of the superior temporal plane for orthogonal penetration of area A1, as determined by pre-implant MRI. Together with a PEEK headpost (to permit painless head restraint), they were secured to the skull with ceramic screws and embedded in dental acrylic. We used MRI compatible materials to allow for post implant imaging. A recovery time of six weeks was allowed before we began behavioral training and data collection.

### Electrophysiology

During the experiments, animals sat in a primate chair in a dark, isolated, electrically shielded, sound-attenuated chamber with head fixed in position, and were monitored with infrared cameras. Neuroelectric activity was obtained using linear array multi-contact electrodes (23 contacts, 100 μm intercontact spacing, Plexon Inc.). The multielectrodes were inserted acutely through guide tube grid inserts, lowered through the dura into the brain, and positioned such that the electrode channels would span all layers of the cortex (Figure 1), which was determined by inspecting the laminar response profile to binaural broadband noise bursts. Neuroelectric signals were impedance matched with a pre-amplifier (10x gain, bandpass dc-10 kHz) situated on the electrode, and after further amplification (500x) they were recorded continuously with a 0.01 – 8000 Hz bandpass digitized with a sampling rate of 44 kHz and precision of 12-bits using Alpha Omega SnR system. The signal was split into the field potential (0.1-300Hz) and MUA (300-5000Hz) range by zero phase shift digital filtering. MUA data was also rectified in order to improve the estimation of firing of the local neuronal ensemble (Legatt et al., 1980). One-dimensional current source density (CSD) profiles were calculated from the local field potential profiles using a three-point formula for the calculation of the second spatial derivative of voltage (Freeman and Nicholson, 1975). The advantage of CSD profiles is that they are not affected by volume conduction like the local field potentials (Kajikawa and Schroeder, 2011, 2015), and they also provide a more direct index of the location, direction, and density of the net transmembrane current flow (Mitzdorf, 1985;Schroeder et al., 1998). At the beginning of each experimental session, after refining the electrode position in the neocortex, we established the best frequency (BF) of the cortical column site using a “suprathreshold” method (Lakatos et al., 2005b; Steinschneider et al., 1995). The method entails presentation of a stimulus train consisting of 100 random order occurrences of a broadband noise burst and pure tone stimuli with frequencies ranging from 353.5 Hz to 32 kHz in half octave steps (duration: 100 ms, r/f time: 5 ms, SOA = 624.5). Auditory stimuli were produced using Tucker Davis Technology’s System III coupled with MF-1 free field speakers.

### Eye Tracking and Saccade Detection

While the animal’s head was immobilized, eye position was monitored at a sampling rate of 500Hz using the Eyelink 1000 Plus system (SR Research Ltd). Vertical and horizontal eye movements were transformed into voltages and simultaneously recorded with the electrophysiological data. Using custom-written functions in Matlab (Mathworks, Natick, MA) we detected saccades if the magnitude of the eye movement (calculated as the length of the eye movement vector) exceeded an arbitrary cut-off value (not measured in distance since our eye tracking system was not calibrated). The cut-off was optimized based on input from several lab members who have experience with manual saccade detection. Saccades had to last for at least 8ms and the minimum distance between saccades (i.e. minimum duration of fixation) had to be at least 120ms.

### Stimulus Presentation

Subjects sat passively in a dark sound attenuated chamber during two conditions: 1) during the presentation of a rhythmic stream of LED flashes which were 25ms in duration and had a stimulus onset asynchrony of 562ms (corresponding to a 1.8Hz repetition rate). We termed this the visual stimulation condition. And 2) in the absence of any stimuli (resting state condition). There was no task in either condition, but the animals were kept alert by interacting with them prior to and after each trial block.

### Data analysis

Data were analysed offline using native and custom-written functions in Matlab. After selective averaging of the CSD and MUA responses to the tones presented in the suprathreshold tonotopy paradigm, recording sites were functionally defined as belonging to A1 based on the sharpness of frequency tuning, the inspection of the tonotopic progression across adjacent sites, and relative sensitivity to pure tones versus broad-band noise of equivalent intensity (Lakatos et al., 2005b; Merzenich and Brugge, 1973; Rauschecker et al., 1997). In the present study only recordings obtained from area A1 were analysed. At the end of each animal’s experimental participation, functional assignment of the recording sites was confirmed histologically (Schroeder et al., 2001).

We utilized the BF-tone related laminar CSD profiles to functionally identify supragranular, granular and infragranular layers in area A1. While the individual layers are not yet possible to exactly differentiate based on electrophysiological criteria, the pial surface, layer 4 and the bottom of layer 6 are easily identifiable – at least in A1 – and using these boundaries and results of prior anatomical studies one can make a pretty accurate estimation (e.g. Fig. 3A-B). To characterize the phase distribution of rhythmic excitability fluctuations as indexed by MUA amplitude at the time of LED flashes or saccades, continuous oscillatory amplitudes and phases of the MUA signal were extracted using two methods: 1) instantaneous phase in single trials was extracted by wavelet decomposition (Morlet wavelet, σ = 6) (Fig. 1F), and 2) the single trial signal was first filtered (in the 1.2-2.4Hz range to extract LED flash related phases, and in the 1-5Hz range to extract saccade timing related phases), and phases were calculated using the Hilbert transform. (Figs. 2 & 3). Inter-trial coherence (ITC) was calculated using the phase values at the time of LED flashes and saccades. ITC ranges from 0 to 1; higher values indicate that the observations (oscillatory phase at a given time-point across LED or saccade events) are clustered more closely around the mean than lower values (phase distribution is biased). Significant phase locking – deviation from uniform (random) phase distribution – was tested with Rayleigh’s uniformity test. The α value was set at 0.05 for all statistical tests, except for the data shown in Figure 3 where the α value was set at 0.01. No corrections for multiple comparisons were applied.

To estimate the strength of MUA entrainment related to saccades vs. LED flashes during a trial block, we calculated LED and saccade related ITC in 10-second-long time windows across the entire trial block (Fig. 4). This yielded 17 LED and approximately 26 saccade related phase values per window. To achieve a relatively good frequency resolution, we used a 1 second stepping of the window, which resulted in a 9 second overlap. Correlation of the “moving” ITC values related to saccades and LED flashes was tested using Spearman’s correlation statistic (Fig. 4).

## Acknowledgments

Support for this work was provided by NIH grants R01DC012947, P50MH109429 and R01MH109289. The authors declare no competing financial interests.

